# A Taxonomically-informed Mass Spectrometry Search Tool for Microbial Metabolomics Data

**DOI:** 10.1101/2023.07.20.549584

**Authors:** Simone Zuffa, Robin Schmid, Anelize Bauermeister, Paulo Wender P. Gomes, Andres M. Caraballo-Rodriguez, Yasin El Abiead, Allegra T. Aron, Emily C. Gentry, Jasmine Zemlin, Michael J. Meehan, Nicole E. Avalon, Robert H. Cichewicz, Ekaterina Buzun, Marvic Carrillo Terrazas, Chia-Yun Hsu, Renee Oles, Adriana Vasquez Ayala, Jiaqi Zhao, Hiutung Chu, Mirte C. M. Kuijpers, Sara L. Jackrel, Fidele Tugizimana, Lerato Pertunia Nephali, Ian A. Dubery, Ntakadzeni Edwin Madala, Eduarda Antunes Moreira, Leticia Veras Costa-Lotufo, Norberto Peporine Lopes, Paula Rezende-Teixeira, Paula C. Jimenez, Bipin Rimal, Andrew D. Patterson, Matthew F. Traxler, Rita de Cassia Pessotti, Daniel Alvarado-Villalobos, Giselle Tamayo-Castillo, Priscila Chaverri, Efrain Escudero-Leyva, Luis-Manuel Quiros-Guerrero, Alexandre Jean Bory, Juliette Joubert, Adriano Rutz, Jean-Luc Wolfender, Pierre-Marie Allard, Andreas Sichert, Sammy Pontrelli, Benjamin S Pullman, Nuno Bandeira, William H. Gerwick, Katia Gindro, Josep Massana-Codina, Berenike C. Wagner, Karl Forchhammer, Daniel Petras, Nicole Aiosa, Neha Garg, Manuel Liebeke, Patric Bourceau, Kyo Bin Kang, Henna Gadhavi, Luiz Pedro Sorio de Carvalho, Mariana Silva dos Santos, Alicia Isabel Pérez-Lorente, Carlos Molina-Santiago, Diego Romero, Raimo Franke, Mark Brönstrup, Arturo Vera Ponce de León, Phillip Byron Pope, Sabina Leanti La Rosa, Giorgia La Barbera, Henrik M. Roager, Martin Frederik Laursen, Fabian Hammerle, Bianka Siewert, Ursula Peintner, Cuauhtemoc Licona-Cassani, Lorena Rodriguez-Orduña, Evelyn Rampler, Felina Hildebrand, Gunda Koellensperger, Harald Schoeny, Katharina Hohenwallner, Lisa Panzenboeck, Rachel Gregor, Ellis Charles O’Neill, Eve Tallulah Roxborough, Jane Odoi, Nicole J. Bale, Su Ding, Jaap S. Sinninghe Damsté, Xueli Li Guan, Jerry J. Cui, Kou-San Ju, Denise Brentan Silva, Fernanda Motta Ribeiro Silva, Gilvan Ferreira da Silva, Hector H. F. Koolen, Carlismari Grundmann, Jason A. Clement, Hosein Mohimani, Kirk Broders, Kerry L. McPhail, Sidnee E. Ober-Singleton, Christopher M. Rath, Daniel McDonald, Rob Knight, Mingxun Wang, Pieter C. Dorrestein

**Author notes:** These authors equally contributed to this work (Simone Zuffa, Robin Schmid, Anelize Bauermeister).

## Abstract

MicrobeMASST, a taxonomically-informed mass spectrometry (MS) search tool, tackles limited microbial metabolite annotation in untargeted metabolomics experiments. Leveraging a curated database of >60,000 microbial monocultures, users can search known and unknown MS/MS spectra and link them to their respective microbial producers via MS/MS fragmentation patterns. Identification of microbial-derived metabolites and relative producers, without *a priori* knowledge, will vastly enhance the understanding of microorganisms’ role in ecology and human health.

## Main

Microorganisms drive the global carbon cycle^1^ and can establish symbiotic relationships with host organisms, influencing their health, aging, and behavior^2–6^. Microbial populations interact with different ecosystems through the alteration of available metabolite pools and the production of specialized small molecules^7,8^. The vast genetic potential of these communities is exemplified by human-associated microorganisms, which encode approximately 100 times more genes than the human genome^9,10^. However, this metabolic potential remains unreflected in modern untargeted metabolomics experiments, where typically <1% of the annotated molecules can be classified as microbial. This problem particularly affects mass spectrometry (MS)-based untargeted metabolomics, a common technique to investigate molecules produced or modified by microorganisms^11^, which famously struggles with spectral annotation of complex biological samples. This is because the majority of spectral reference libraries are biased towards commercially available or otherwise accessible standards of primary metabolites, drugs, or industrial chemicals. Even when metabolites are annotated, extensive literature searches are required to understand whether these molecules have microbial origins and to identify the respective microbial producers. Public databases, such as KEGG^12^, MiMeDB^13^, NPAtlas^14^, and LOTUS^15^, can assist in this interpretation, but they are mostly limited to well-established, largely genome-inferred, metabolic models or to fully characterized and published molecular structures. Additionally, while targeted metabolomics efforts aimed at interrogating the gut microbiome mechanistically have been developed^16^, these focus only on relatively few commercially-available microbial molecules. Hence, the majority of the microbial chemical space remains unknown, despite the continuous expansion of MS reference libraries. To fill this gap, we have developed microbeMASST (https://masst.gnps2.org/microbemasst/), a search tool that leverages public MS repository data to identify the microbial origin of known and unknown metabolites and map them to their microbial producers.

MicrobeMASST is a community-sourced tool that works within the GNPS^17^ ecosystem. Users can search tandem MS (MS/MS) spectra obtained from their experiments against MS/MS spectra previously detected in other extracts of bacterial, fungal, or archaeal monocultures. No other available resource or tool allows linking uncharacterized MS/MS spectra to characterized microorganisms. The microbeMASST reference database of monocultures has been generated through years of community contributions and metadata curation, and it contains microorganisms isolated from plants, soils, oceans, lakes, fish, terrestrial animals, and humans (**Figure 1a**). All available microorganisms are categorized according to the NCBI taxonomy^18^ at different taxonomic resolution (*i*.*e*. species, genus, family, *etc*.) or mapped to the closest taxonomically accurate level, if no NCBI ID was available at the time of database creation. As of June 2023, microbeMASST includes 60,781 liquid chromatography (LC)-MS/MS files, comprising >100 million MS/MS spectra, mapped to 541 strains, 1,336 species, 539 genera, 264 families, 109 orders, 41 classes, and 16 phyla from the three domains of life: Bacteria, Archaea, and Eukaryota (**Figure 1b**). Differently from MASST^19^, which uses a precomputed network of ∼110 million MS/MS spectra to enable spectral searching, microbeMASST is based on the newly introduced Fast Search Tool (https://fasst.gnps2.org/fastsearch/)^20^. This tool, originally designed for proteomics, drastically improves search speed by several orders of magnitude by indexing all the MS/MS spectra present in GNPS/MassIVE and restricting the search space to the user input parameters. Because of this, search results are returned within seconds as opposed to 20 min per search or 24-48 hours for modification tolerant searches in the original implementation of MASST. Additionally, microbeMASST leverages the pre-curated file-associated metadata to aggregate results into taxonomic trees. This represents a major enhancement over MASST, where users have to manually inspect result tables and contextualize them, making interpretations tedious.

**Figure 1.**
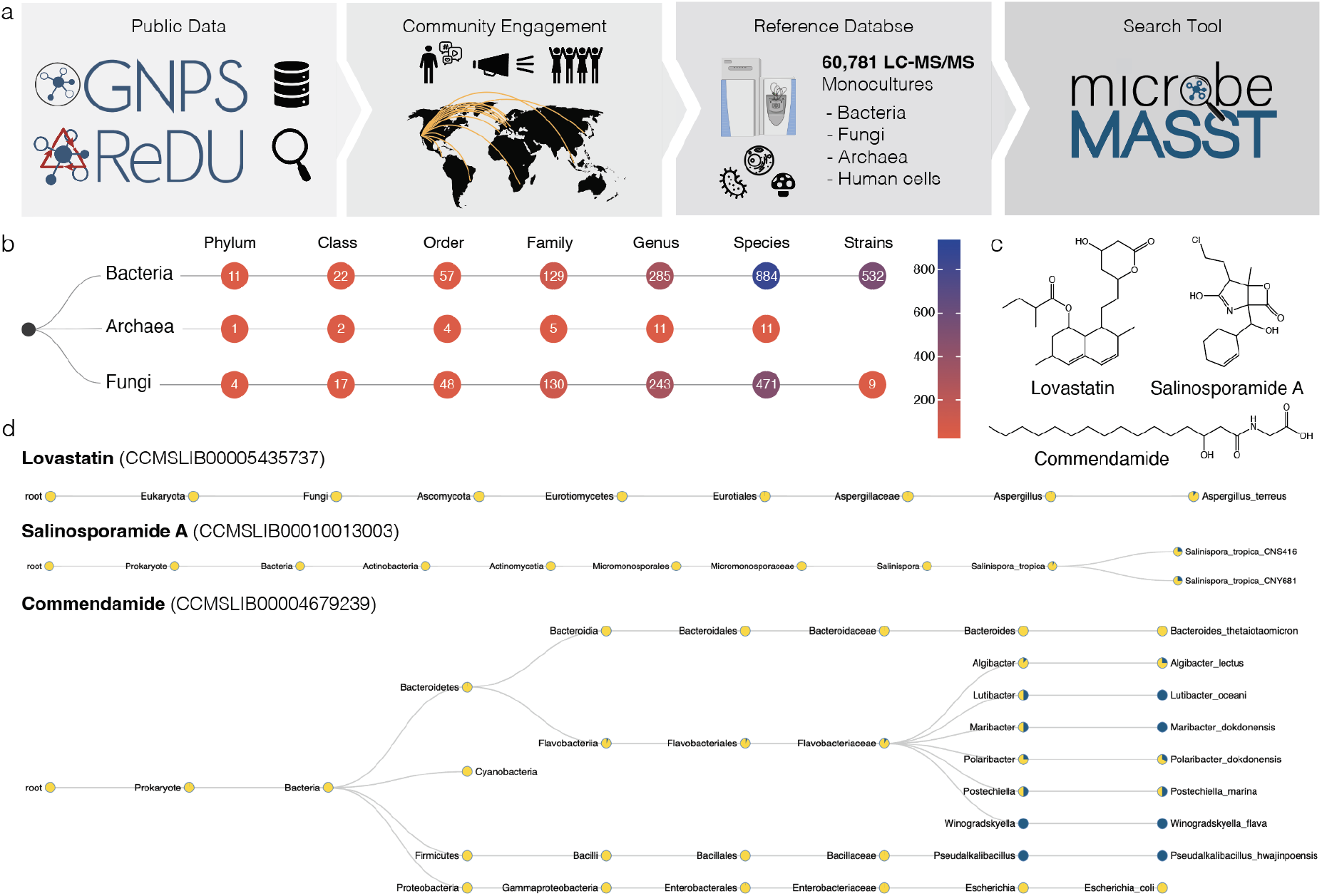
The microbeMASST search tool and reference database. **a**) Community contributions of data and knowledge to GNPS^17^, ReDU^23^, and MassIVE from 2014 to 2022 were used to generate the microbeMASST reference database. Additionally, a public invitation to deposit data in June 2022 resulted in the further deposition of LC-MS/MS files from 25 different laboratories from 15 different countries across the globe, leading to the curation of a total of 60,781 LC-MS/MS files of microbial monoculture extracts. **b**) MicrobeMASST comprises 1,858 unique lineages, across three different domains of life, mapped to 541 unique strains, 1,336 species, 539 genera, 264 families, 109 orders, 41 classes, and 16 phyla. **c**) Examples of medically-relevant small molecules known to be produced by bacteria or fungi. Lovastatin, a cholesterol lowering drug originally isolated from *Aspergillus* genus^24^, salinosporamide A, a Phase III candidate to treat glioblastoma produced by *Salinispora tropica*^*25*^, and commendamide, a human G-protein–coupled receptor agonist^26^. **d**) MicrobeMASST search outputs of the three different molecules of interest confirm that they were exclusively found in monocultures of the only known producers. Pie charts display the proportion of MS/MS matches found in the deposited reference database. Blue indicates a match with a monoculture, while yellow represents a nonmatch. Searches were performed using MS/MS spectra deposited in the GNPS reference library: lovastatin (<u>CCMSLIB00005435737</u>), salinosporamide A (<u>CCMSLIB00010013003</u>), and commendamide (<u>CCMSLIB00004679239</u>).

In microbeMASST, users can search MS/MS spectra using a Universal Spectrum Identifier (USI)^21^ or by inputting a precursor ion mass and its spectral fragmentation pattern (**Supplementary Figure 1**). Analogue search can also be enabled to discover molecules related to the MS/MS spectrum of interest across the taxonomic tree^17,19,22^. The microbeMASST web app displays query results in interactive taxonomic trees, which can be downloaded as HTML files. Nodes in the trees represent specific taxa and display rich information, such as taxon scientific name, NCBI taxonomic ID, number of deposited samples, number of found MS/MS matches, and proportion of found matches, which is also visualized through pie charts. Information for an MS/MS match in a particular taxon is propagated upstream through its lineage. The reactive interface of microbeMASST enables filtering of the tree to specific taxonomic levels or to a minimum number of matches observed per taxon. Additionally, three data tables are generated, linking the search job to other resources in the GNPS/MassIVE ecosystem. Each MS/MS query is searched against the public MS/MS reference library of GNPS (587,213 MS/MS spectra, June 2023). Annotations to such reference compounds are listed under the ‘Library matches’ tab (**Supplementary Figure 2a**). The ‘Datasets matches’ tab contains information on the matching scans, displaying scientific name, NCBI taxonomic ID and taxonomic rank, number of matching fragment ions, and modified cosine score together with a link to a mirror plot visualization (**Supplementary Figure 2b**). Finally, the ‘Taxa matches’ tab informs on how many matches were found per taxon and number of samples available for that taxon (**Supplementary Figure 2c**). Quality controls (QCs) and blank samples (n=2,902) present in the reference datasets of microbeMASST have been retained to provide information on possible contaminants and media components. Additionally, data from human cell line cultures (n=1,199) have been included to enable assessment of whether molecules can be produced by both human hosts and microorganisms.

Search results for lovastatin, salinosporamide A, and commendamide MS/MS spectra highlight how microbeMASST can correctly connect microbial molecules to their known producers (**Figure 1c**). In the case of lovastatin, a clinically-used cholesterol-lowering drug originally isolated from *Aspergillus terreus*^*24*^, spectral matches were unique to the genus *Aspergillus* (**Figure 1d**). The MS/MS spectrum for salinosporamide A, a Phase III candidate to treat glioblastoma^27^, only matched two strains of *Salinispora tropica* (**Figure 1d**), the only known producer^25^. Commendamide, first observed in cultures of *Bacteroides vulgatus* (recently reclassified as *Phocaeicola vulgatus*), is a G-protein–coupled receptor agonist^26^. Surprisingly it had many matches to several bacterial cultures, including in Flavobacteriaceae (*Algibacter, Lutibacter, Maribacter, Polaribacter, Postechiella*, and *Winogradskyella*) and *Bacteroides* cultures (**Figure 1d**). Additional examples include searches of mevastatin, arylomycin A4, yersiniabactin, promicroferrioxamine, and the microbial bile acid conjugates^28–30^ glutamate-cholic acid (Glu-CA) and glutamate-deoxycholic acid (Glu-DCA) (**Supplementary Figure 3**). Mevastatin, another cholesterol-lowering drug originally isolated from *Penicillium citrinum*^*31*^, was only found in samples classified as fungi. The antibiotic arylomycin A4 was observed in different *Streptomyces* species and it was originally isolated from *Streptomyces* sp. Tue 6075 in 2002^32^. Yersiniabactin, a siderophore originally isolated from *Yersinia pestis*^*33*^, whose monoculture is not yet present in the reference database of microbeMASST, was observed in *Escherichia coli* and *Klebsiella* species, consistent with previous observations^34,35^. Promicroferrioxamine, another siderophore, was observed to match *Micromonospora chokoriensis* and *Streptomyces* species. This molecule was originally isolated from an uncharacterized *Promicromonosporaceae* isolate^36^. The MS/MS spectrum of the gut microbiota-derived Glu-CA, an amidated tri-hydroxylated bile acid, was most frequently observed in cultures of *Bifidobacterium* species, while Glu-DCA was found only in one *Bifidobacterium* strain but also in two *Enterococcus* and *Clostridium* species. None of the aforementioned molecules were found in cultured human cell lines, highlighting the ability of microbeMASST to distinguish MS/MS spectra of molecules that can be exclusively produced by either bacteria or fungi. It is important to acknowledge that MS/MS data generally do not differentiate stereoisomers, but it can nevertheless provide crucial information on molecular families.

MicrobeMASST can be also used to extract microbial information from mass spectrometry-based metabolomics studies without any *a priori* knowledge. To illustrate this, we reprocessed an untargeted metabolomics study comparing germ-free (GF) mice to those harboring microbial communities, also known as specific pathogen-free (SPF) mice^29^ (**Figure 2a**). We extracted 10,047 consensus MS/MS spectra uniquely present in SPF mice and queried them with microbeMASST. A total of 3,262 MS/MS spectra were found to have a microbial match. Of these, 837 were also found in human cell lines and for this reason were removed from further analysis. Among the remaining 2,425 MS/MS spectra, 1,673 were exclusively found in bacteria, 95 in fungi, and 657 in both (**Supplementary Figure 4**). These MS/MS spectra were then processed with SIRIUS^37^ and CANOPUS^38^ to tentatively annotate the metabolites and identify their chemical classes. A file containing all these spectra of interest can be explored and downloaded as .mgf format from GNPS (**see Methods**). To further validate the microbial origin of these MS/MS spectra, we assessed their overlap with data acquired from a different study comparing SPF mice treated with a cocktail of antibiotics to untreated controls^40^. Interestingly, 621 MS/MS spectra were also found in this second dataset and 512 were only present in animals not treated with antibiotics (**Figure 2b**). The distribution of these spectra and their classes across bacterial phyla was visualized using an UpSet plot^39^ (**Figure 2c**). Notably, the majority of the spectra classified as terpenoids were commonly observed across phyla while amino acids and peptides appeared to be more phylum specific. Of these 512 spectra, 23% had a level 2 or 3 annotation^41^, matching against the GNPS reference libraries (**Supplementary Table 1**). These included the recently described amidated microbial bile acids^19,28–30,42–47^, free bile acids originating from the hydrolysis of host derived taurine bile acid conjugates^48^, keto bile acids formed via microbial oxidation of alcohols^29^, *N*-acyl-lipids belonging to a similar class of metabolites as commendamide^26^ (a microbial *N*-acyl lipid), di- and tri-peptides seen in microbial digestion of proteins^49^, and soyasapogenol, a byproduct of the microbial digestion of complex saccharides from dietary soyasaponins^29^. Part of the remaining unannotated spectra can be identified as chemical modifications of the above annotated microbial metabolites through spectral similarity obtained from molecular networking (**Supplementary Figure 5**). Based on literature information, the list of annotated MS/MS spectra contained a small number of metabolites traditionally considered to be non-microbial in origin. One interpretation of this finding is that microorganisms are capable of producing metabolites previously described to only be made by mammalian hosts. Notable examples include serotonin^50^, γ-aminobutyric acid (GABA)^51^, and the glycocholic acid^42,52–54^, with microorganisms often being the primary producers of these metabolites in the gut. Additionally, an alternative hypothesis is that microorganisms can also selectively stimulate the production of host metabolites. Other limitations regarding annotations are discussed in **Methods**. To assess if the observations from the mouse models translate to humans, we searched and found that 455 out of the 512 MS/MS spectra of interest matched to public human data (**Figure 2d**). Interestingly, these spectra were found in both healthy individuals and individuals affected by different health states, including type II diabetes, inflammatory bowel disease (IBD), Alzehimer’s diseases and other conditions. These spectra were most commonly found in stool samples (n=110,973 MS/MS matches) followed by blood, breast milk, and the oral cavity as well as other organs including the brain, skin, vagina, and biofluids, such as cerebrospinal fluid and urine (**Figure 2e**). These findings support the concept that a significant number of microbial metabolites reach and influence distant organs in the human body^55^.

**Figure 2.**
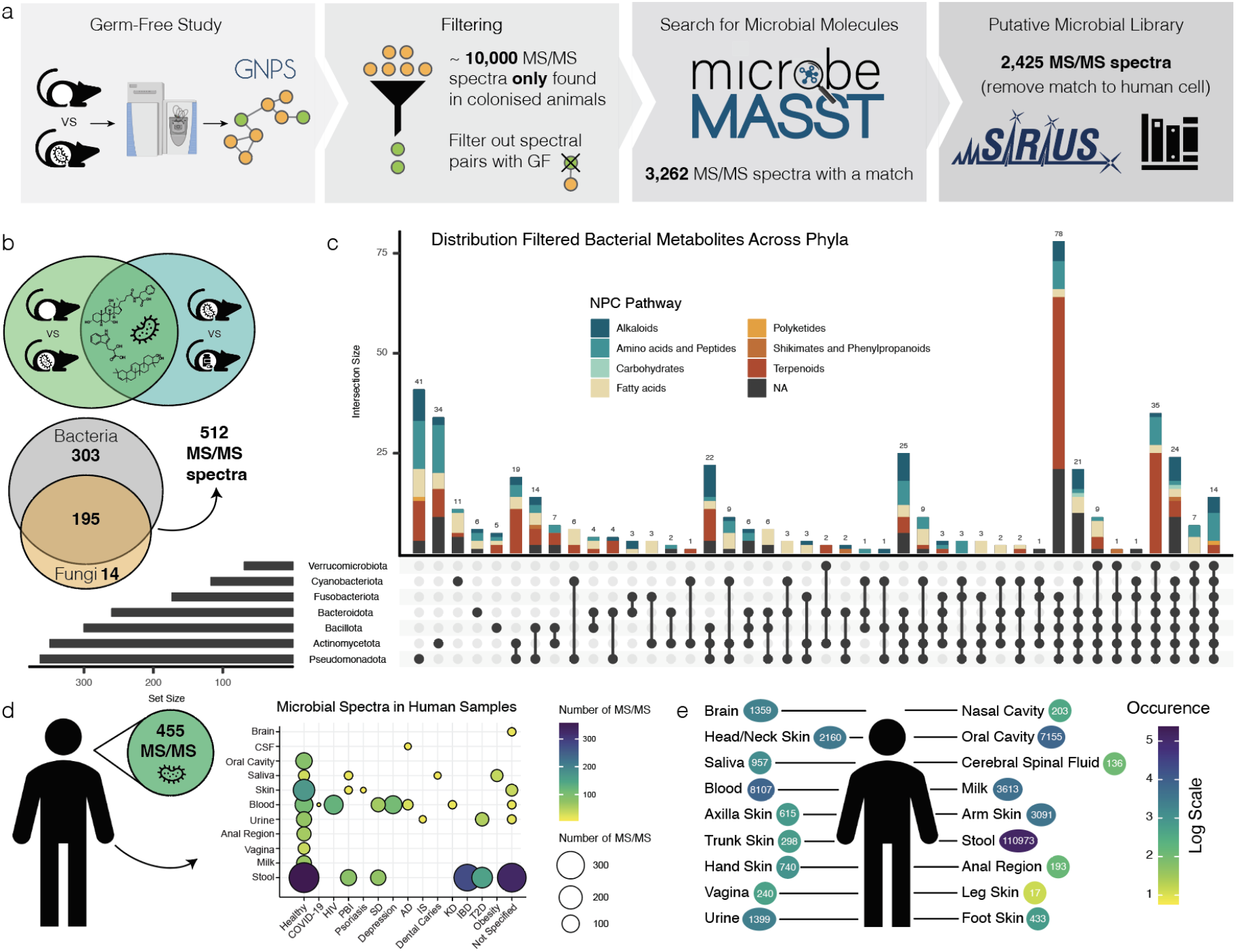
MicrobeMASST can identify microbial MS/MS spectra within mouse and human datasets. **a)** Workflow to extract microbial MS/MS spectra from biochemical profiles of 29 different tissues and biofluids of SPF mice that are not observed in GF mice^29^. The MS/MS spectra unique to SPF mice (10,047) were searched with microbeMASST. A total of 3,262 MS/MS spectra had a match; those MS/MS also matching human cell lines were removed, leaving a total of 2,425 putative microbial MS/MS spectra (**see Methods** to download .mgf file). **b**) The presence of the 2,425 MS/MS spectra was evaluated in an additional animal study looking at antibiotics usage^40^. A total of 512 MS/MS spectra, out of the 621 overlapping, was exclusively found in animals not receiving antibiotics. **c**) UpSet plot of the distribution of the detected MS/MS spectra (512) across bacterial phyla. Terpenoids were more commonly observed across phyla while amino acids and peptides appeared to be more phylum specific **d**) The 512 MS/MS spectra were searched in human datasets and 455 were found to have a match. These MS/MS spectra were present in both healthy individuals and individuals affected by different diseases. **e**) Most of the MS/MS spectra (n=411) matched fecal samples (n=110,973 matches), followed by blood, oral cavity, breast milk, urine, and several other organs. Abbreviations: CSF, cerebral spinal fluid; HIV, human immunodeficiency virus; PBI, primary bacterial infectious disease; SD, sleep disorder; AD, Alzheimer’s disease; IS, ischemic stroke; KD, Kawasaki disease; IBD, inflammatory bowel disease; T2D, type II diabetes.

We anticipate microbeMASST will be a key resource to enhance understanding of the role of microbial metabolites across a wide range of ecosystems, including oceans, plants, soils, insects, animals, and humans. This expanding resource will enable the scientific community to gain valuable taxonomic and functional insights into diverse microbial populations. The mass spectrometry community will play a key role in the evolution of this tool in the future through the continued deposition of data associated with novel microbial monocultures and the expansion of spectral reference libraries. Moreover, microbeMASST holds potential for various applications, ranging from aquaculture and agriculture to biotechnology and the study of microbial-mediated human health conditions. By harnessing the power of public data, we can unlock new opportunities for advancements in multiple fields and deepen our understanding of the intricate relationships between microorganisms and their ecosystems.

## Acknowledgments

This work was carried out through the collaborative microbial metabolite center, which is supported by the National Institutes of Health (NIH) grant U24DK133658, the Alzheimer’s gut project U19AG063744 and BBSRC-NSF award 2152526. We thank Travis Adkins and Lisa McCormick from the USDA ARS Culture Collection for their assistance selecting and providing microbial strains used in this research. This project was supported in part by the U.S. Department of Agriculture, Agricultural Research Service. AMCR and HM were supported by the NIH grant 1DP2GM137413. Research reported in this publication was supported in part by the National Center for Complementary and Integrative Health of the NIH under award number F32AT011475 to NEA. KBK is supported by National Research Foundation of Korea (NRF) grants funded by the Korean Government (MSIT) (NRF-2020R1C1C1004046, 2022R1A5A2021216, and 2022M3H9A2096191). BS is supported by the Austrian Science Fund (FWF) P31915 and KH is financed by the FWF research group program (grant FG3). DP is supported by the German Research Foundation (DFG) CMFI Cluster of Excellence (EXC 2124) and Collaborative Research Center CellMap (TRR 261). AB, EAM, COG, PCJ, LVCL, NPL are supported by The São Paulo Research Foundation - FAPESP #2018/24865-4, #2019/03008-9, #2020/06430-0, #2022/12654-4, #2015/17177-6, #2020/02207-5, #2021/10603-0) and CNPq. CLC received financial support from StrainBiotech and the FEMSA Biotechnology Centre from Tecnológico de Monterrey. LRO received a scholarship from the Mexican National Council of Science and Technology (CONACYT). PCT and WHG are supported by NIH R01 GM107550. NG is supported by the NSF CAREER Award #2047235. NS, SD and JSSD are supported by ERC Horizon 2020 (grant agreement No 694569) and by a Spinoza award to JSSD from NWO. BW is supported by CMFI Cluster of Excellence (EXC 2124). MS is supported by DZIF (Grant no TTU 09.722). AVPDL, SLLR, and PP are supported by ERA-Net Cofund project BlueBio (grant agreement no. 311913), Research Council of Norway (300846). HMR, MFL and GLB are supported by Novo Nordisk Foundation (grant NNF19OC0056246; PRIMA—toward Personalized dietary Recommendations based on the Interaction between diet, Microbiome and Abiotic conditions in the gut). HMR is supported by the Independent Research Fund Denmark (MOTILITY; grant no. 0171-00006B). ECON is supported by the Nottingham Research Fellowship. AIPL is supported by FPU (FPU19/00289). MFT and RdCP were supported by R35GM12889.CMS is supported by Juan de la Cierva-Incorporación (IJC2018-036923-I) and Proyectos dirigidos por jóvenes investigadores de la Universidad de Málaga (B1-2021_21). DR is supported by Plan Nacional de I+D+i of the Ministerio de Ciencia e Innovación (PID2019-107724GB-I00), and Junta de Andalucía (P20_00479). XLG is supported by Nanyang Assistant Professorship. KSJ is supported by NIH R01 GM137135. AS is supported by EMBO fellowship ALTF 996-2021. SP is supported by ETH Zurich Career Seed Fellowship. RG is supported by the Simons Foundation Postdoctoral Fellowship in Marine Microbial Ecology. HC is supported by NIH R01AI167860 and CIFAR. MCT is supported by T32 DK007202 (NIDDK), the National Academies of Sciences, Engineering and Medicine through the Predoctoral Fellowship of the Ford Foundation, and the Howard Hughes Medical Institute (HHMI) Graduate Fellowships grant (GT15123). EEL is supported by VI-Universidad de Costa Rica, Grants numbers C1604 and C0469. PE is supported by VI-Universidad de Costa Rica, Grants numbers C1604 and C0469; U.S. National Science Foundation DEB-1638976. DAV and GTC are supported by VI-Universidad de Costa Rica, Grants numbers C1604 and C0469. KF is supported by CMFI Cluster of Excellence (EXC 2124). HHFK and GFS are supported by Fundação de Amparo à Pesquisa do Estado do Amazonas (FAPEAM). DBS is supported by Fundação de Apoio ao Desenvolvimento do Ensino, Ciência e Tecnologia do Estado de Mato Grosso do Sul - FUNDECT (process number: 71/032.390/2022, FUNDECT number: 311/2022). KLM is supported by NIH / 1R01GM132649. PMA is supported by a swissuniversities Open Research Data grant. MCMK and SLJ are supported by NIH R35GM142938. ADP is supported by NIH U01 DK119702 and S10 OD021750. ATA was supported by the Betty and Gordon Moore Foundation. ML and PB are supported by the Max Planck Society. Omnia Group Ltd. is duly acknowledged for microbial cultures. BSP and NB were partially supported by NIH 1R01LM013115 and NSF ABI 1759980. RK is supported by NIH DP1AT010885. LPN is supported by Omnia Group Ltd. Shimadzu South Africa Ltd. is gratefully thanked for the analytical support. We thank James MacRae, head of The Metabolomics STP at the Francis Crick Institute for his guidance.

## Author contributions (names given as initials)

SZ, RS, AB, MW, PCD - Conceptualized the method

RS, SZ, MW - Developed microbeMASST

PCD, SZ, AB, PWPG, AMCR, YEA, ATA, ECG, JZ, MJM, NEA, RHC, EB, MCT, CYH, RO, AVA, JZ, HC, MCMK, SJ, FT, LPN, NEM, IAD, EAM, LVCL, NPL, PRT, PCJ, BR, ADP, MFT, RCP, GTC, PC, EEL, DAV, LMQG, JLW, AS, SP, JJ, KG, JMC, PMA, BCW, KF, DP, NA, NG, ML, PB, KBK, AGG, HG, LPSC, MSS, AIPL, CMS, DR, RF, MB, AVPL, PBP, SLLR, GLB, MFL, HMR, AR, BS, FH, AJB, CLC, LRO, ER, FH, GK, HS, KH, LP, RG, ECON, ETR, JO, NJB, SD, XLG, JJC, KSJ, DBS, FMRS, GFS, HHFK, JAC, HM, KB, KLM, SOS, CMR, and RK Contributed data and curated metadata

SZ - Generated taxonomic tree and performed analyses

RS - Developed the tree visualizer for enriched ontologies and output summaries

BSP, NB - Developed the FASST algorithm

MW - Developed the Fast Search Tool API

SZ, RS, PCD - Tested microbeMASST

SZ, RS, PCD - Wrote the manuscript

All authors reviewed the manuscript

## Data and code availability

Data used to generate the reference database of microbeMASST are publicly available at GNPS/MassIVE (https://massive.ucsd.edu/). A list with all the accession numbers (MassIVE IDs) of the studies used to generate this tool is available in **Supplementary Table 2**. Interactive examples of the MS/MS queries illustrated in **Figure 1d** and **Supplementary Figure 3** can be generated, visualized, and downloaded from the microbeMASST website (https://masst.gnps2.org/microbemasst/). Known molecules already present in the GNPS library (https://library.gnps2.org/) were used to facilitate interpretation and confirm that specific bacterial and fungal molecules were exclusively observed in the respective monocultures.

- Lovastatin - <u>CCMSLIB00005435737</u>
- Salinosporamide A - <u>CCMSLIB00010013003</u>
- Commendamide - <u>CCMSLIB00004679239</u>
- Mevastatin - <u>CCMSLIB00005435644</u>
- Arylomycin A4 - <u>CCMSLIB00000075066</u>
- Yersiniabactin - <u>CCMSLIB00005435750</u>
- Promicroferrioxamine - <u>CCMSLIB00005716848</u>
- Glutamate-cholic acid (Glu-CA) - <u>CCMSLIB00006582258</u>
- Glutamate-deoxycholic acid (Glu-DCA) - <u>CCMSLIB00006582092</u>

Data used to extract MS/MS spectra exclusively present in colonized (SPF) mice is publicly available in GNPS/MassIVE under the accession number <u>MSV000079949</u>. Data used to validate and assess antibiotics effect on microbial MS/MS spectra of interest is available under the accession number <u>MSV000080918</u>. List of datasets with data acquired from human biosamples that presented matches to the putative microbial MS/MS spectra of interest is available in **Supplementary Table 3**.

The microbeMASST code to query spectra, create interactive trees, and analyze results is available under open source license on GitHub (https://github.com/robinschmid/microbe_masst). Code used to generate the microbeMASST web interface can be accessed on GitHub (https://github.com/mwang87/GNPS_MASST). Code used to perform the analysis and generate the figures present in the manuscript can be downloaded from GitHub (https://github.com/simonezuffa/Manuscript_microbeMASST)

### Data collection and harmonization

Data deposited in GNPS/MassIVE was investigated manually and systematically, using ReDU^23^ (https://redu.ucsd.edu/), to extract all the publicly available MS/MS files (.mzML or .mzXML formats) acquired from monocultures of bacteria, fungi, archaea, and human cell lines. Only monocultures were included in this search tool to unequivocally associate the production of the detected metabolites to each specific taxon. A total of 60,781 files from 537 different GNPS/MassIVE datasets were selected to be used as a reference database of microbeMASST (**Supplementary Table 2**). These comprise files deposited in response to our call to the scientific community. Between May and July 2022, 25 different research groups deposited 65 distinct datasets in GNPS/MassIVE, comprising a total of 3,142 unique LC-MS/MS files. This represented a 5.45% increase in publicly available MS/MS data acquired from monocultures in just two months. To qualify as a contributor and be credited as one of the authors, researchers had to deposit high resolution LC-MS/MS data acquired either in positive or negative ionization modes from monocultures of either bacteria, fungi, or achaea. Harmonization of the acquired data and metadata represented a challenge. The NCBI taxonomic database is constantly expanding and evolving and ReDU latest updated (December 2021) does not accommodate the latest deposited taxa. For this reason, an additional metadata file (microbeMASST_metadata_*massiveID*) was generated specifically for the microbeMASST project and uploaded to the respective GNPS/MassIVE datasets deposited by the collaborators, if the ReDU workflow failed. All the collected information was finally aggregate in one single .csv file (microbe_masst_table.csv) that can be found on GitHub, which contains: 1) Full MassIVE path of each sample, 2) Filename of each sample reported as its MassIVE ID/file name to avoid presence of duplicated names, 3) MassIVE ID, 4) Taxonomic name of the isolate as reported by the author submitting the associated metadata, 5) Alternative taxonomic name if the provided taxonomic name was incorrect or not present in NCBI, 6) Associated NCBI ID to the taxonomic name or the alternative taxonomic name, when present, 7) Definition if the taxonomic ID was automatically assigned or manually curated, and information if 8) ReDU metadata is available for that specific file and if the file correspond to a 9) Blank or 10) Quality control (QC) rather than an unique biological sample.

Unique taxonomic names and NCBI IDs were extracted from the metadata associated with the selected samples. When metadata was not available and multiple species of bacteria or fungi were present in the same dataset, samples were generically classified as bacteria or fungi. Concordance between taxonomic names and NCBI IDs was checked by blasting taxonomic names against NCBI (https://www.ncbi.nlm.nih.gov/Taxonomy/TaxIdentifier/tax_identifier.cgi) to obtains respective NCBI IDs and updated taxonomic names. Results were manually investigated and missing IDs were recovered using the NCBI browser (https://www.ncbi.nlm.nih.gov/Taxonomy/Browser/wwwtax.cgi). If the taxonomic name was not found in NCBI, most likely because it was not deposited yet, the NCBI of the closest taxon was retrieved and used. For example, the strain *Staphylococcus aureus* CM05 was unavailable in NCBI and was curated to the species *Staphylococcus aureus* instead.

### Taxonomic tree generation

The microbeMASST taxonomic tree was generated using both R 4.2.2 (R Foundation for Statistical Computing) and Python 3.10 (Python Software Foundation). In R, the microbeMASST table was filtered and only unique NCBI IDs were retained (n = 1,834). The classification function from the ‘taxize’ package (v 0.9.100) was used to retrieve the full lineage of each NCBI ID^56^. Main taxonomic ranks (kingdom to strain) plus subgenus, subspecies, and varieties were kept in order to obtain taxonomic trees with a similar number of nodes per lineage. The list of NCBI IDs of all lineages was then imported in Python, where the ETE3 toolkit was used to generate a comprehensive taxonomic tree based on the provided NCBI IDs^57^. The generated Newick tree was then converted into JSON format and information such as taxonomic rank, number of samples available was added. Additionally, children nodes for blanks and QCs were created in order to be visualized in the same tree, if observed.

### MASST query

MicrobeMASST web application was built using Dash and Flask open source libraries for Python (https://github.com/mwang87/GNPS_MASST/blob/master/dash_microbemasst.py). The web app can receive as inputs either a USI^21^ or an MS/MS spectrum (fragment ions and their intensities). Additionally, batch searches can be performed using a customisable python script that can read either a .tsv file containing a list of USIs or a single .mgf spectra file. Through the manuscript we showcase how we were able to search for more than 10,000 MS/MS spectra contained in a single .mgf file (approximately 2 hours run time). After receiving input information, microbeMASST leverages the Fast Search Tool (https://fasst.gnps2.org/fastsearch/) API and the sample-specific associated metadata to generate taxonomic trees. Fast searches are based on indexing all the MS/MS spectra present in GNPS/MassIVE according to the mass and intensity of their precursor ions and then restricting the search to only a set of relevant spectra that have a greater chance to achieve a high spectral similarity (modified cosine score) to the MS/MS of interest. Searches are customizable and default settings are the following: precursor and fragment ion mass tolerances = 0.05, minimum cosine score threshold = 0.7, minimum number of matching fragment ions = 3, and analog search = False. The JSON file of the microbeMASST taxonomic tree is then filtered according to the results and converted into a D3 JavaScript object that can be visualized as an HTML file.

### Applications

We envision microbeMASST having several applications. First, we showcase how researchers can investigate single MS/MS spectra, using the web interface, and obtain matching results if their known or unknown MS/MS spectrum was previously observed in any of the microbial monocultures present in the microbeMASST database. Nine small molecules of interest were investigated using MS/MS spectra already deposited in the GNPS reference library (see **Data and code availability**). Second, we show how microbeMASST can be leveraged to mine for known or unknown microbial metabolites in MS studies. To test this hypothesis, we reprocessed an LC-MS/MS dataset acquired from GF and SPF mice^29^. A comprehensive molecular network was generated (https://gnps.ucsd.edu/ProteoSAFe/status.jsp?task=893fd89b52dc4c07a292485404f97723). From the obtained job, the qiime2 artifact (qiime2_table.qza), the .mgf file (METABOLOMICS-SNETS-V2-893fd89b-download_clustered_spectra-main.mgf) containing all the captured MS/MS spectra, and the annotation table (METABOLOMICS-SNETS-V2-893fd89b-view_all_annotations_DB-main.tsv) were extracted. The .qza file first converted into a .biom file and then .tsv using QIIME2^58^ to extract the feature table. This was then imported in R where only spectra present in tissues and biofluids of SPF animals were retained (n = 10,047). To add an extra layer of filtering, all MS/MS spectra that had an edge (cosine similarity > 0.7) and a delta parent ion mass +/- 0.02 Da with MS/MS spectra present in GF animals were removed (spectral pairs information is contained in networkedges_selfloop). All the MS/MS spectra were then run in batch using a custom python script of microbeMASST (processing time: ∼2 hours, Apple M2 Max, 64GB RAM) to obtain microbial matches. Matching and filtered MS/MS spectra (n = 2.425) were aggregated into a single .mgf file that can be downloaded from GNPS (https://gnps.ucsd.edu/ProteoSAFe/status.jsp?task=aecd30b9febd43bd8f57b88598a05553). Compound class of each MS/MS spectrum, with parent ion mass < 850 Da, was predicted with SIRIUS^37^ and CANOPUS^38^. The 2,425 MS/MS spectra were then searched against the MSV000080918 dataset containing animals treated or not with antibiotics^40^. Matching and filtered MS/MS spectra (n = 512) were aggregated into a single .mgf file that can be downloaded from GNPS (https://gnps.ucsd.edu/ProteoSAFe/status.jsp?task=c33855fc32c948049331e9730189d5c1). A list of the spectra with putative chemical class classification is available in **Supplementary Table 1**. Venn diagrams and UpSet plots were generated in R using ‘VennDiagram 1.7.3’, ‘UpSetR 1.4.0’, and ‘ComplexUpset 1.3.3’. Finally, the 512 MS/MS spectra were searched in batch against the GNPS public repository to observe if they were detected in human datasets (**Supplementary Table 3**)

### Technical limitations

Analysis of the results should be considered with these limitations in mind. Molecule detection in microbeMASST is dependent on the availability of specific substrates in the reference monocultures. If the culture lacks the necessary substrates (or any other culture condition) to produce a certain molecule, this molecule will not be detected. Nevertheless, if related substrates are present then a different but related molecule may be produced instead, which can be detected using the analog search. To address this problem, it is crucial for the community to continue to gather data from as many diverse experimental conditions as possible to capture the full range of metabolites produced by different microorganisms. This will help in building the most comprehensive reference database that encompasses diverse microbial metabolic profiles. Isomers and stereoisomers, which are molecules with the same molecular formula but different structural arrangements, often exhibit similar MS/MS spectra. This means that their fragmentation patterns may not provide enough information to distinguish them. Differences in extraction conditions and instrument settings can lead to variations in the obtained MS/MS spectra. For example, the intensity of precursor ions used for fragmentation can impact the resulting spectra. If the precursor ion intensity is low, the fragmented spectrum may lack ions that are present in spectra obtained from high-intensity precursor ions. This may result in “data leakage”, as the MS/MS spectrum may be missing ions, and leading to the two molecules not being recognized as the same molecule. To partially overcome this more permissive settings can be created. The majority of the data deposited in public repositories, GNPS included, and used in microbeMASST were acquired using positive ionization mode, which limits the observation of molecules that cannot be ionized in positive mode. This means that certain metabolites may be underrepresented or not detected at all. The continuous curation of the microbeMASST reference database involves adding more diverse data in terms of ionization modes to improve the coverage of metabolites. Taxonomic tree was generated using associated NCBI IDs provided by the community. Specimen assignment prior to metabolomic analysis can not be checked by microbeMASST. The majority of the deposited data do not have associated genetic information and even if available, it was not used to build taxonomic tree. Thus, specimen mis-identification is possible. By addressing these challenges and continuously curating the reference database with more comprehensive and diverse data, microbeMASST coverage can be expanded to provide valuable insights into the role of microbiota and to facilitate our understanding of microbial metabolism in diverse ecosystems.

## Supplementary Figures

**Supplementary Figure 1.**
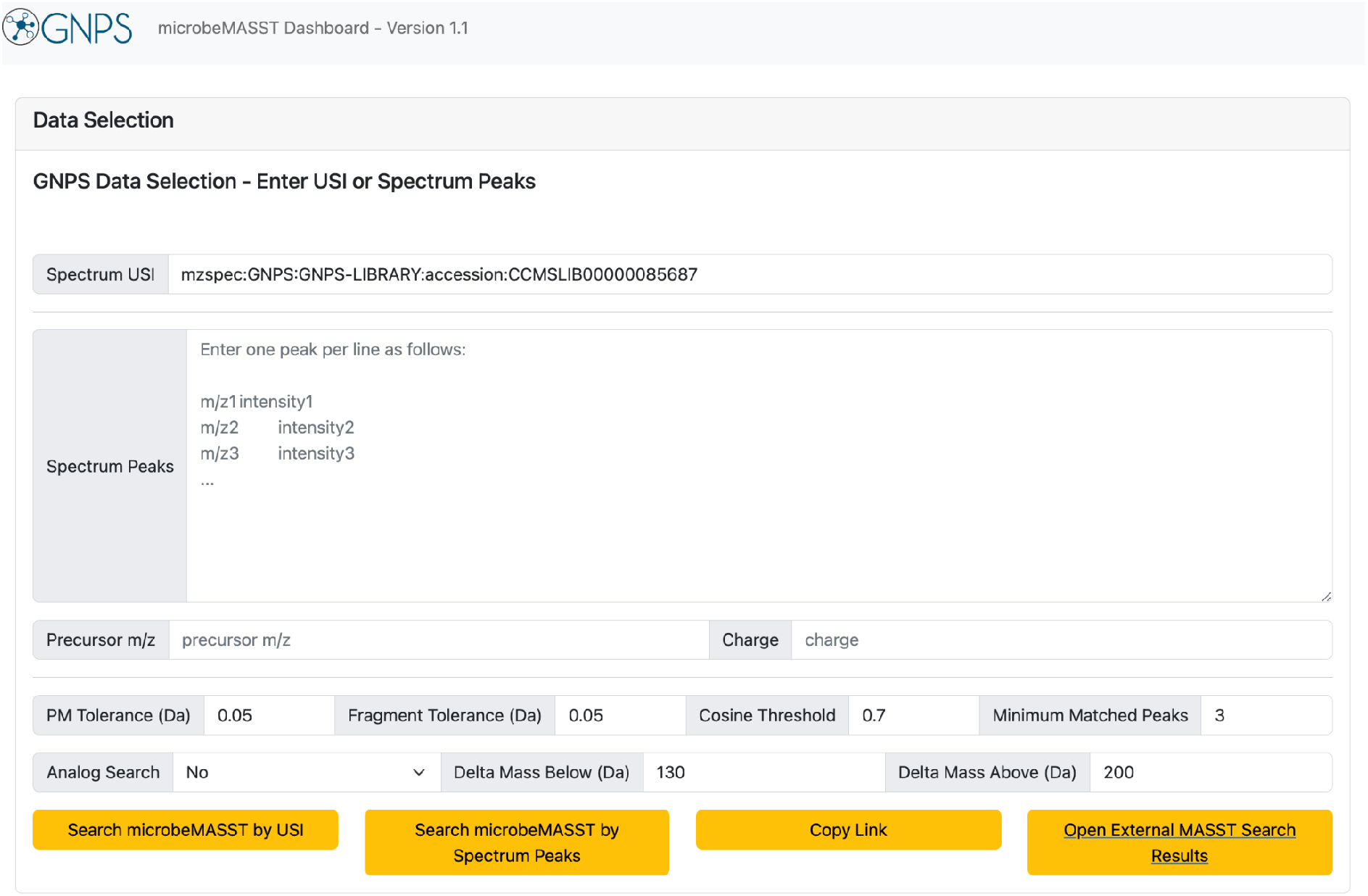
microbeMASST web app. Users can access the microbeMASST web app at https://masst.gnps2.org/microbemasst/. They can search for MS/MS spectrum against the microbeMASST reference database by either providing a USI in ‘Spectrum USI’ or by inputting fragment ions and their intensities and the precursor mass in ‘Spectrum Peak’ and ‘Precursor m/z’ respectively. Search parameters, such as parent ion and fragment ions tolerances, cosine threshold, and minimum matching peaks can be modified. Analog search can also be enabled. Finally, to submit a search query users have just to click either on ‘Search microbeMASST by USI’ or ‘Search microbeMASST by Spectrum Peak’ based on the information that they have provided. Search jobs can be easily shared by clicking on ‘Copy Link’.

**Supplementary Figure 2.**
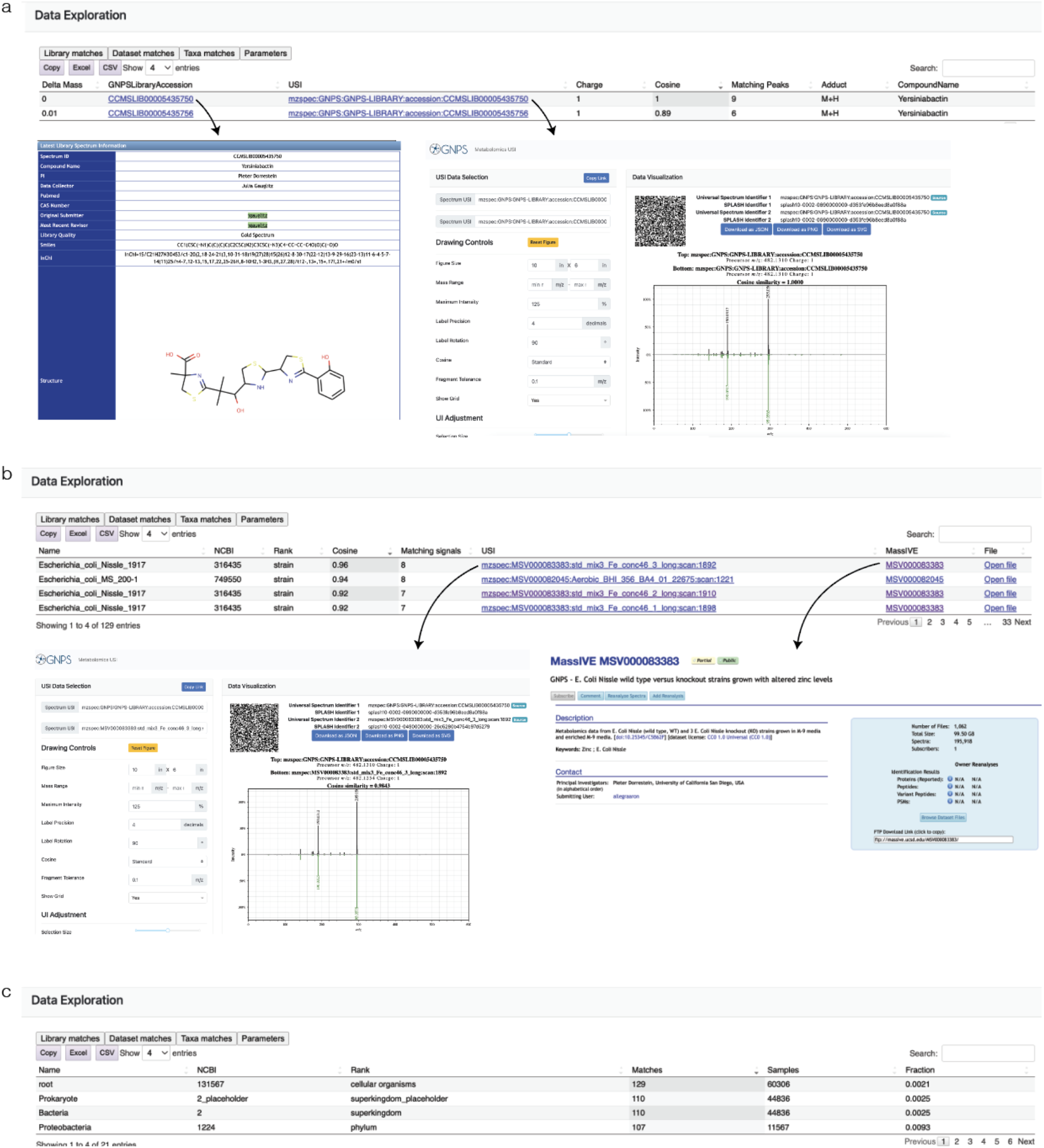
Complementary output of microbeMASST. **a)** The queried MS/MS spectrum is searched against the GNPS libraries and, if matches are found, possible annotations are returned. Information on cosine similarity and number of matching peaks is provided and users can explore the associated GNPS Library Spectrum page and inspect mirror plots. **b)** Information of matching scans in the sample from the different taxa is provided. In addition to cosine similarity score, matching fragments, and the possibility to inspect mirror plots, users can retrieve the MassIVE ascension number of the project together with contact information on who deposited the data. **c)** Number of matches for each taxonomic level are returned. Users can observe how many samples for that specific taxon are part of microbeMASST and see how broadly the molecule is distributed within it (fraction).

**Supplementary Figure 3.**
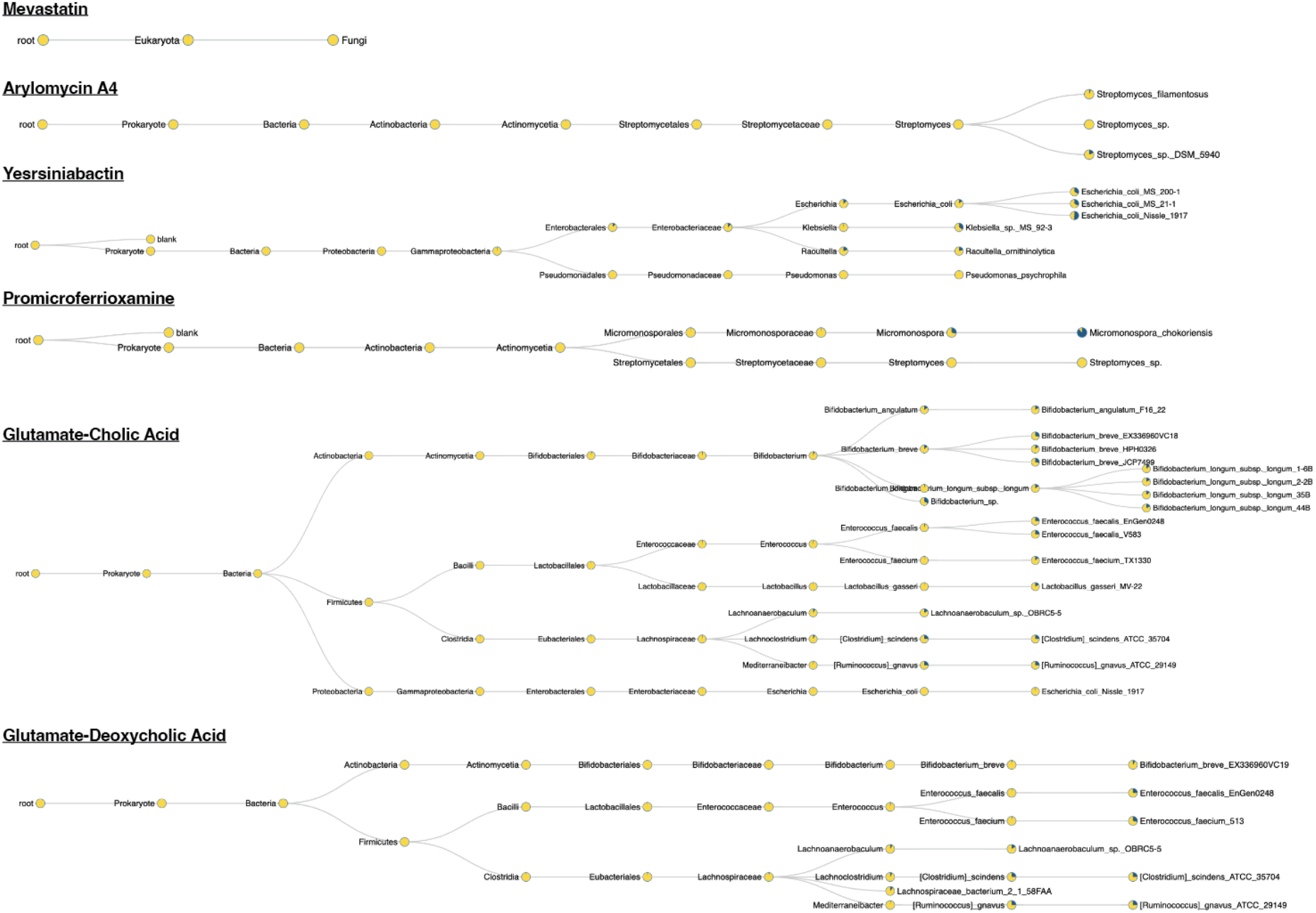
Examples of microbeMASST outputs. Additional examples of results obtained from searching from <u>mevastatin</u>, <u>arylomycin A4</u>, <u>yersiniabactin</u>, <u>promicroferrioxamine</u>, <u>glutamate-cholic acid</u>, and <u>glutamate deoxycholic acid</u>. In all cases the molecules were found in the monocultures of known producers and not in human cell lines, confirming that microbeMASST can be used to search for microbial-derived molecules.

**Supplementary Figure 4.**
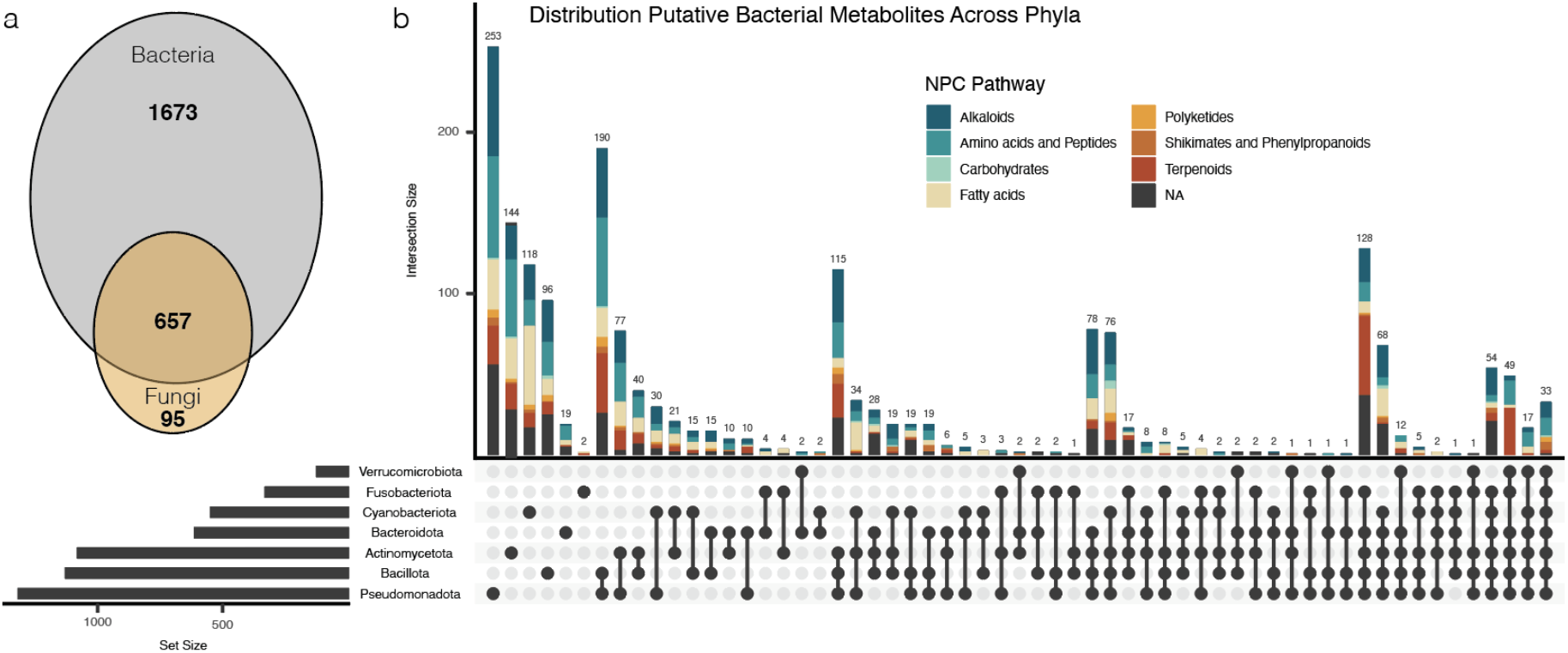
Distribution putative microbial metabolites across phyla. a) Of the 2,425 MS/MS spectra that had a match exclusively to microbial monocultures, 1,673 were found only in bacteria, 95 in fungi, and 657 in both. b) Chemical classes were then predicted using SIRIUS and CANOPUS and their distribution across the different phyla was visualized using an Upset plot.

**Supplementary Figure 5.**
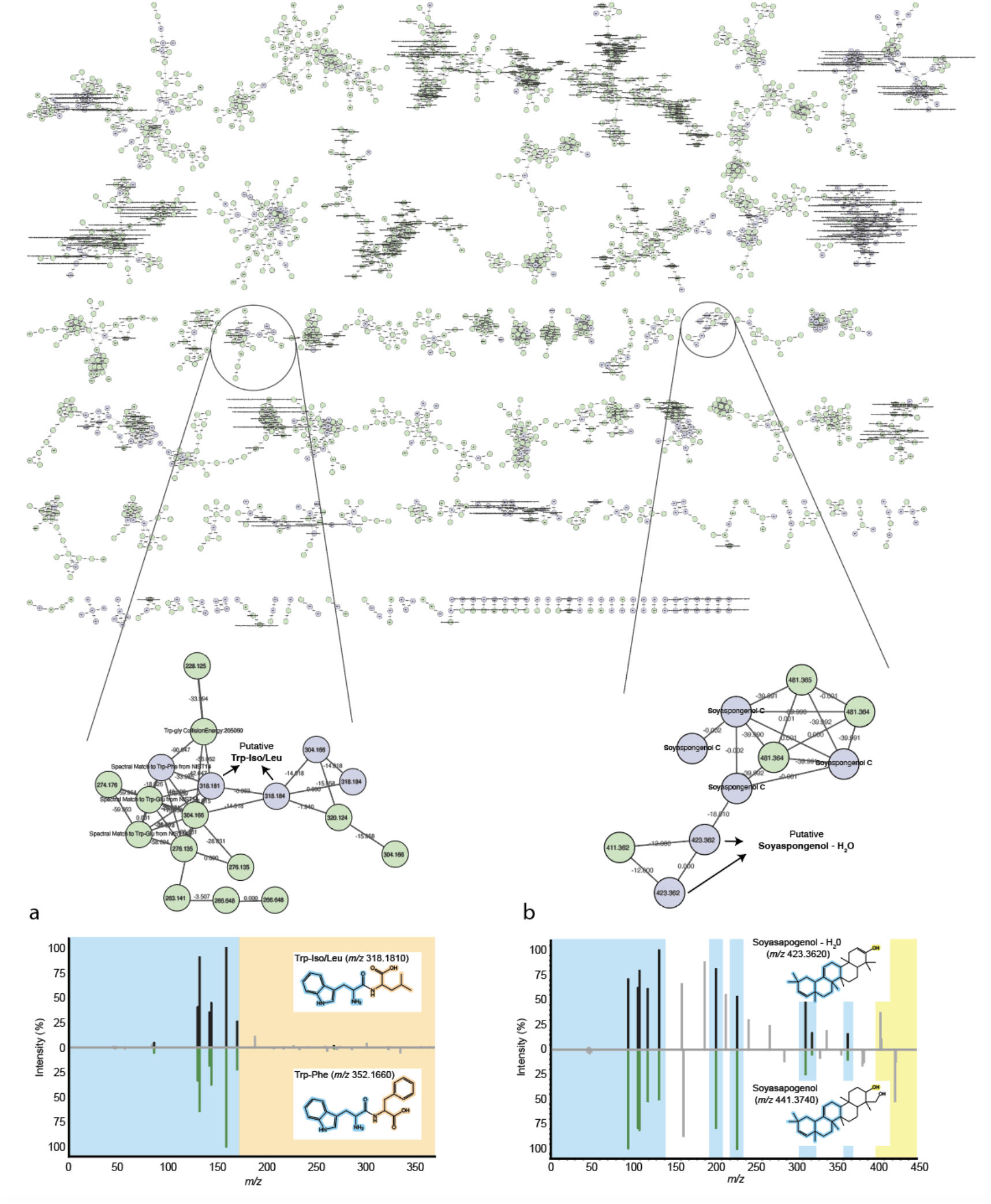
Contextualization of the molecular network. The selected 512 MS/MS spectra were mapped back to the full molecular network generated from the data acquired from GF and SPF mice (MSV000079949). Only molecular families containing at least one of the MS/MS spectra of interest were retained and used to generate putative annotations of the unannotated spectra of interest. Each node represents a spectrum (ion), light blue indicates MS/MS of interest while green indicates spectra that were not retained in the performed downstream analysis. Within each node the *m*/*z* of the precursor ion is indicated, while on each edge the *m*/*z* difference between two connected nodes is reported. a) Ions with precursor mass 318.181 and directly connected to a spectrum annotated as the dipeptide Trp-Phe (*m*/*z* 352.144) can be tentatively annotated as Trp-Iso/Leu. b) Ions with precursor mass 423.362 connected to Soyasapogenol C, which can represent a soyasapogenol molecule with a loss of water (delta mass 18.01). Abbreviations: Trp, tryptophan; Phe, phenylalanine; Iso, isoleucine; Leu, leucine.

## References

1. Jansson, J. K. & Hofmockel, K. S. Soil microbiomes and climate change. Nat. Rev. Microbiol. 18, 35–46 (2020).

2. Malard, F., Dore, J., Gaugler, B. & Mohty, M. Introduction to host microbiome symbiosis in health and disease. Mucosal Immunol. 14, 547–554 (2021).

3. Fan, Y. & Pedersen, O. Gut microbiota in human metabolic health and disease. Nat. Rev. Microbiol. 19, 55–71 (2021).

4. López-Otín, C., Blasco, M. A., Partridge, L., Serrano, M. & Kroemer, G. Hallmarks of aging: An expanding universe. Cell 186, 243–278 (2023).

5. Radjabzadeh, D. et al.. Gut microbiome-wide association study of depressive symptoms. Nat. Commun. 13, 7128 (2022).

6. Morais, L. H., Schreiber, H. L., 4th & Mazmanian, S. K. The gut microbiota-brain axis in behaviour and brain disorders. Nat. Rev. Microbiol. 19, 241–255 (2021).

7. Milshteyn, A., Colosimo, D. A. & Brady, S. F. Accessing Bioactive Natural Products from the Human Microbiome. Cell Host Microbe 23, 725–736 (2018).

8. Paoli, L. et al.. Biosynthetic potential of the global ocean microbiome. Nature 607, 111–118 (2022).

9. Grice, E. A. & Segre, J. A. The human microbiome: our second genome. Annu. Rev. Genomics Hum. Genet. 13, 151–170 (2012).

10. Tierney, B. T. et al.. The Landscape of Genetic Content in the Gut and Oral Human Microbiome. Cell Host Microbe 26, 283–295.e8 (2019).

11. Bauermeister, A., Mannochio-Russo, H., Costa-Lotufo, L. V., Jarmusch, A. K. & Dorrestein, P. C. Mass spectrometry-based metabolomics in microbiome investigations. Nat. Rev. Microbiol. 20, 143–160 (2022).

12. Kanehisa, M. & Goto, S. KEGG: kyoto encyclopedia of genes and genomes. Nucleic Acids Res. 28, 27–30 (2000).

13. Wishart, D. S. et al.. MiMeDB: the Human Microbial Metabolome Database. Nucleic Acids Res. 51, D611–D620 (2023).

14. van Santen, J. A. et al.. The Natural Products Atlas 2.0: a database of microbially-derived natural products. Nucleic Acids Res. 50, D1317–D1323 (2022).

15. Rutz, A. et al.. The LOTUS initiative for open knowledge management in natural products research. Elife 11, (2022).

16. Han, S. et al.. A metabolomics pipeline for the mechanistic interrogation of the gut microbiome. Nature 595, 415–420 (2021).

17. Wang, M. et al.. Sharing and community curation of mass spectrometry data with Global Natural Products Social Molecular Networking. Nat. Biotechnol. 34, 828–837 (2016).

18. Schoch, C. L. et al.. NCBI Taxonomy: a comprehensive update on curation, resources and tools. Database 2020, (2020).

19. Wang, M. et al.. Mass spectrometry searches using MASST. Nat. Biotechnol. 38, 23–26 (2020).

20. Batsoyol, N., Pullman, B., Wang, M., Bandeira, N. & Swanson, S. P-massive: a real-time search engine for a multi-terabyte mass spectrometry database. in Proceedings of the International Conference on High Performance Computing, Networking, Storage and Analysis 1–15 (IEEE Press, 2022).

21. Deutsch, E. W. et al.. Universal Spectrum Identifier for mass spectra. Nat. Methods 18, 768–770 (2021).

22. Watrous, J. et al.. Mass spectral molecular networking of living microbial colonies. Proc. Natl. Acad. Sci. U. S. A. 109, E1743–52 (2012).

23. Jarmusch, A. K. et al.. ReDU: a framework to find and reanalyze public mass spectrometry data. Nat. Methods 17, 901–904 (2020).

24. Alberts, A. W. et al.. Mevinolin: a highly potent competitive inhibitor of hydroxymethylglutaryl-coenzyme A reductase and a cholesterol-lowering agent. Proc. Natl. Acad. Sci. U. S. A. 77, 3957–3961 (1980).

25. Fenical, W. et al.. Discovery and development of the anticancer agent salinosporamide A (NPI-0052). Bioorg. Med. Chem. 17, 2175–2180 (2009).

26. Cohen, L. J. et al.. Functional metagenomic discovery of bacterial effectors in the human microbiome and isolation of commendamide, a GPCR G2A/132 agonist. Proc. Natl. Acad. Sci. U. S. A. 112, E4825–34 (2015).

27. CTG Labs - NCBI. https://beta.clinicaltrials.gov/study/NCT03345095?distance=50&intr=NPI-0052&rank=9.

28. Dorrestein, P. et al.. A synthesis-based reverse metabolomics approach for the discovery of chemical structures from humans and animals. (2021) doi:10.21203/rs.3.rs-820302/v1.

29. Quinn, R. A. et al.. Global chemical effects of the microbiome include new bile-acid conjugations. Nature 579, 123–129 (2020).

30. Patterson, A. et al.. Bile Acids Are Substrates for Amine N-Acyl Transferase Activity by Bile Salt Hydrolase. (2022) doi:10.21203/rs.3.rs-2050120/v1.

31. Endo, A. The origin of the statins. 2004. Atheroscler. Suppl. 5, 125–130 (2004).

32. JUDITH Schimana, KLAUS Gebhardt Alexand R. HÖLtzel, DIETMAR G. Schmid, RODERICH SÜSsmuth, Johannes MÜLler, RÜDiger Pukall, HANS-PETER FIEDLER. Arylomycins A and B, New Biaryl-bridged Lipopeptide Antibiotics Produced by Streptomyces sp. Tü 6075. The Journal of Antibiotics (2002).

33. Drechsel, H. et al.. Structure elucidation of yersiniabactin, a siderophore from highly virulent Yersinia strains. Liebigs Ann.: Org. Bioorg. Chem. 1995, 1727–1733 (1995).

34. Schubert, S., Picard, B., Gouriou, S., Heesemann, J. & Denamur, E. Yersinia high-pathogenicity island contributes to virulence in Escherichia coli causing extraintestinal infections. Infect. Immun. 70, 5335–5337 (2002).

35. Lawlor, M. S., O’connor, C. & Miller, V. L. Yersiniabactin is a virulence factor for Klebsiella pneumoniae during pulmonary infection. Infect. Immun. 75, 1463–1472 (2007).

36. Yang, Y.-L. et al.. Connecting chemotypes and phenotypes of cultured marine microbial assemblages by imaging mass spectrometry. Angew. Chem. Int. Ed Engl. 50, 5839–5842 (2011).

37. Dührkop, K. et al.. SIRIUS 4: a rapid tool for turning tandem mass spectra into metabolite structure information. Nat. Methods 16, 299–302 (2019).

38. Dührkop, K. et al.. Systematic classification of unknown metabolites using high-resolution fragmentation mass spectra. Nat. Biotechnol. 39, 462–471 (2021).

39. Lex, A., Gehlenborg, N., Strobelt, H., Vuillemot, R. & Pfister, H. UpSet: Visualization of Intersecting Sets. IEEE Trans. Vis. Comput. Graph. 20, 1983–1992 (2014).

40. Shalapour, S. et al.. Inflammation-induced IgA+ cells dismantle anti-liver cancer immunity. Nature 551, 340–345 (2017).

41. Sumner, L. W. et al.. Proposed minimum reporting standards for chemical analysis Chemical Analysis Working Group (CAWG) Metabolomics Standards Initiative (MSI). Metabolomics 3, 211–221 (2007).

42. Lucas, L. N. et al.. Dominant Bacterial Phyla from the Human Gut Show Widespread Ability To Transform and Conjugate Bile Acids. mSystems e0080521 (2021).

43. Hoffmann, M. A. et al.. High-confidence structural annotation of metabolites absent from spectral libraries. Nat. Biotechnol. 40, 411–421 (2022).

44. Guzior, D. et al.. Bile salt hydrolase/aminoacyltransferase shapes the microbiome. (2022) doi:10.21203/rs.3.rs-2050406/v1.

45. Foley, M. H. et al.. Bile salt hydrolases shape the bile acid landscape and restrict Clostridioides difficile growth in the murine gut. Nat Microbiol 8, 611–628 (2023).

46. Folz, J. et al.. Human metabolome variation along the upper intestinal tract. Nat Metab (2023) doi:10.1038/s42255-023-00777-z.

47. Shalon, D. et al.. Profiling the human intestinal environment under physiological conditions. Nature 617, 581–591 (2023).

48. Yao, L. et al.. A selective gut bacterial bile salt hydrolase alters host metabolism. Elife 7, (2018).

49. Bartlett, A. & Kleiner, M. Dietary protein and the intestinal microbiota: An understudied relationship. iScience 25, (2022).

50. Yano, J. M. et al.. Indigenous bacteria from the gut microbiota regulate host serotonin biosynthesis. Cell 161, 264–276 (2015).

51. Strandwitz, P. et al.. GABA-modulating bacteria of the human gut microbiota. Nat Microbiol 4, 396–403 (2019).

52. Maneerat, S., Nitoda, T., Kanzaki, H. & Kawai, F. Bile acids are new products of a marine bacterium, Myroides sp. strain SM1. Appl. Microbiol. Biotechnol. 67, 679–683 (2005).

53. Kim, D. et al.. Biosynthesis of bile acids in a variety of marine bacterial taxa. J. Microbiol. Biotechnol. 17, 403–407 (2007).

54. Ohashi, K., Miyagawa, Y., Nakamura, Y. & Shibuya, H. Bioproduction of bile acids and the glycine conjugates by Penicillium fungus. J. Nat. Med. 62, 83–86 (2008).

55. Lai, Y. et al.. High-coverage metabolomics uncovers microbiota-driven biochemical landscape of interorgan transport and gut-brain communication in mice. Nat. Commun. 12, 6000 (2021).

56. Chamberlain, S. A. & Szöcs, E. taxize: taxonomic search and retrieval in R. F1000Res. 2, 191 (2013).

57. Huerta-Cepas, J., Serra, F. & Bork, P. ETE 3: Reconstruction, Analysis, and Visualization of Phylogenomic Data. Mol. Biol. Evol. 33, 1635–1638 (2016).

58. Bolyen, E. et al.. Reproducible, interactive, scalable and extensible microbiome data science using QIIME 2. Nat. Biotechnol. 37, 852–857 (2019).

